# Paternal age in rhesus macaques is positively associated with germline mutation accumulation but not with measures of offspring sociability

**DOI:** 10.1101/706705

**Authors:** Richard J. Wang, Gregg W.C. Thomas, Muthuswamy Raveendran, R. Alan Harris, Harshavardhan Doddapaneni, Donna M. Muzny, John P. Capitanio, Predrag Radivojac, Jeffrey Rogers, Matthew W. Hahn

## Abstract

Mutation is the ultimate source of all genetic novelty and the cause of heritable genetic disorders. Mutational burden has been linked to complex disease, including neurodevelopmental disorders such as schizophrenia and autism. The rate of mutation is a fundamental genomic parameter and direct estimates of this parameter have been enabled by accurate comparisons of whole-genome sequences between parents and offspring. Studies in humans have revealed that the paternal age at conception explains most of the variation in mutation rate: each additional year of paternal age in humans leads to approximately 1.5 additional mutations inherited by the child. Here, we present an estimate of the *de novo* mutation rate in the rhesus macaque (*Macaca mulatta*) using whole-genome sequence data from 32 individuals in four large pedigrees. We estimated an average mutation rate of 0.58 × 10^-8^ per base pair per generation (at an average parental age of 7.5 years), much lower than found in direct estimates from great apes (including human, chimpanzee, and gorilla). As in humans, older macaque fathers transmit more mutations to their offspring, approximately 1.5 extra mutations per year in our probands. Mutations at CpG sites accounted for 24% of all observed point mutations. We found that the rate of mutation accumulation after puberty is similar between macaques and humans, but that a smaller number of mutations accumulate before puberty in macaques. We additionally investigated the role of paternal age on offspring sociability, a proxy for normal neurodevelopment. In 203 male macaques studied in large social groups, we found no relationship between paternal age and multiple measures of social function. Our findings are consistent with the hypothesis that the increased risk of neurodevelopmental disorders with paternal age in primates is not primarily due to *de novo* mutations.

## Introduction

Paternal age at conception is the single strongest predictor of the number of *de novo* mutations that a human will inherit. Studies show that children will inherit approximately 1.5 additional mutations per year of paternal age, and that the average father contributes new mutations at a rate that is three to four times greater than the mother per year (Kong et al. 2012; Besenbacher et al. 2015; Francioli et al. 2015; Jónsson et al. 2017). This male bias in the number of *de novo* mutations has been attributed to the continuous nature of spermatogenesis, which results in the accumulation of replication errors in the male germline (Crow 2000). As new mutations are responsible for the incidence of spontaneous genetic disorders, this pattern has considerable implications for human health. The male bias in mutation rate has also been observed in other primates (Venn et al. 2014; Thomas et al. 2018), making it an important feature in models of mutation rate evolution (Thomas and Hahn 2014; Amster and Sella 2016; Moorjani et al. 2016; Besenbacher et al. 2019).

Children conceived by older fathers are at greater risk for numerous adverse developmental outcomes (Crow 2000). These include a greater risk of certain genetic disorders due to new mutations inherited at single genes, as with Apert syndrome, Marfan syndrome, and achondroplasia (Glaser et al. 2003; Green et al. 2010; Goriely et al. 2013). Evidence from epidemiological studies also suggests a relationship between advanced paternal age and complex neurodevelopmental disorders: an increased risk of schizophrenia, autism, and bipolar disorder have all been associated with advanced paternal age (Sipos et al. 2004; Reichenberg et al. 2006; Durkin et al. 2008; Frans et al. 2008; Menezes et al. 2010; Lee et al. 2011). The mechanisms underlying these epidemiological associations with paternal age have not been conclusively determined. A central question lies in whether the risk inherited from older fathers comes from genetic predisposition or *de novo* mutations. In the *de novo* model, neurological disorders are highly polygenic and result from the additive effects of a rising mutational burden (Malaspina et al. 2002; Hultman et al. 2011; Kong et al. 2012; Ronemus et al. 2014). The competing predisposition hypothesis attributes the epidemiological association to underlying pre-existing genetic factors that may actually contribute to delayed reproduction in males (Gratten et al. 2016; Janecka et al. 2017). For example, a genetic correlation between psychiatric disorders and delayed fatherhood could explain the association seen with advanced paternal age.

Primate models provide a powerful experimental system for investigating the relationship between the increasing number of *de novo* mutations and the increased risk of developmental outcomes with paternal age. Direct estimation of the mutation rate by tracking the appearance of new germline mutations across generations has become possible with advances in the availability and economy of next-generation sequencing technologies. While there are some technical concerns in sampling and sequencing of rare *de novo* events (Ségurel et al. 2014; Scally 2016), there are few barriers to estimating mutation rates in any species. Along with estimates of *de novo* mutations, testing this relationship also requires information on neurodevelopmental maturation and expressed behavior in non-human systems. Although such data are much harder to collect, several long-term studies of captive primate populations have tracked multiple aspects of neurological and social abilities. In the important model system, rhesus macaque (*Macaca mulatta*), researchers have found high levels of social intelligence (Capitanio 1999; Sclafani et al. 2016). As a consequence, the rhesus monkey has become a model for studying schizophrenia (Gil-da-Costa et al. 2013) and autism spectrum disorder (Bauman and Schumann 2018; Parker et al. 2018).

Studying the mutation rate in non-human primates also allows us to address fundamental questions concerning its evolution. Because the average parental age at conception in the rhesus macaque is much lower than in human, any process that leads to a relationship between parental age and the number of germline mutations will also lead to a lower per-generation mutation rate in macaques. Clearly, a smaller number of germline cell divisions from spermatogenesis in male macaque parents predicts a smaller number of replication-dependent errors (Thomas and Hahn 2014). Other differences in spermatogenesis between species, such as variation in seminiferous epithelial cycle length and the fraction of apoptotic spermatocytes, could also lead to changes in the total number of cell divisions in the male germline (Amster and Sella 2016; Gao et al. 2016; Scally 2016). The observation of a significant, though smaller, maternal age effect on the human mutation rate (Goldmann et al. 2016; Wong et al. 2016; Jónsson et al. 2017) raises the possibility that a substantial portion of germline mutations may not originate from replicative errors during spermatogenesis (Gao et al. 2019; Sasani et al. 2019). Nonetheless, any relationship between non-replicative mutations and parental age would contribute to the evolution of mutation rate between species due to differences in the age of reproduction. Understanding the contribution of each of these processes to possible differences in the per-generation mutation rate between species—as well as their effects on variation in the substitution rate—is a goal of recent mutation studies of non-human primates (Thomas et al. 2018; Besenbacher et al. 2019).

Here, we performed whole-genome sequencing on 32 rhesus macaques from four multiple-generation pedigrees maintained at the California National Primate Research Center to identify *de novo* mutations. We also analyzed behavioral data in 203 male macaques from the same population to examine links between paternal age at conception and behavioral metrics of sociability, including those linked to autism.

## Results

### The number of mutations inherited increases with paternal age in rhesus monkeys

We sequenced 32 individuals from 4 three-generation families of rhesus monkeys (Fig. S1) to 40× average coverage using Illumina short-read sequencing. These families contained 14 trios with different offspring, from which we identified 307 *de novo* single nucleotide mutations (Table 1). After applying stringent quality filters, comparing two genotype-calling pipelines, and controlling for the callability of sites across the genome (Methods), we estimated an average mutation rate of 0.58 × 10^-8^ per base pair (bp), per generation for parents with an average age of 7.5 years (7.8 paternal, 7.1 maternal).

**Table 1.**
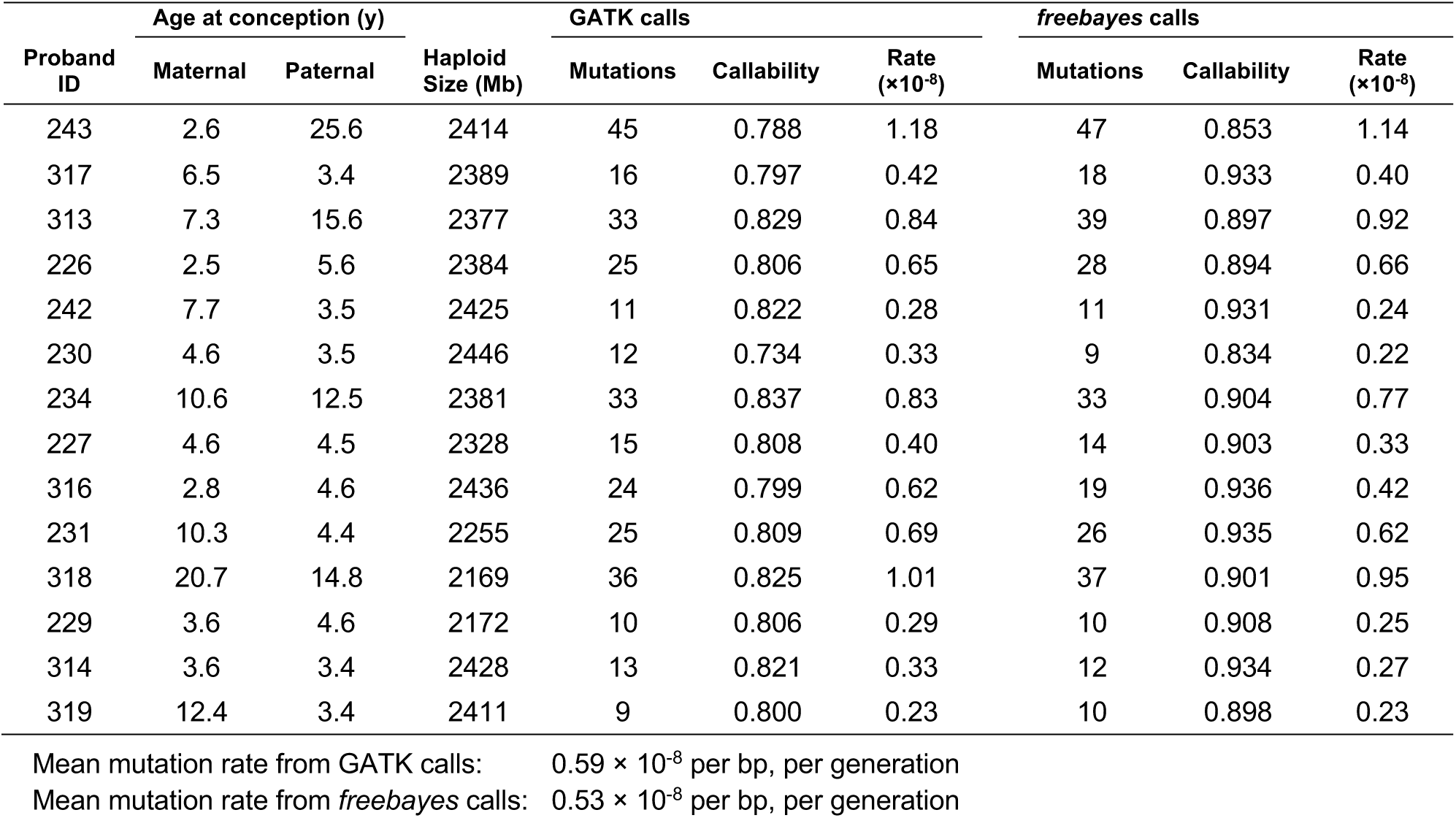
Mutation count and rate by trio

| Proband ID | Age at conception (y) |  | Haploid Size (Mb) | GATK calls |  |  | *freebayes* calls |  |  |
| --- | --- | --- | --- | --- | --- | --- | --- | --- | --- |
| | Maternal | Paternal | | Mutations | Callability | Rate ($\times 10^{-8}$) | Mutations | Callability | Rate ($\times 10^{-8}$) |
| 243 | 2.6 | 25.6 | 2414 | 45 | 0.788 | 1.18 | 47 | 0.853 | 1.14 |
| 317 | 6.5 | 3.4 | 2389 | 16 | 0.797 | 0.42 | 18 | 0.933 | 0.40 |
| 313 | 7.3 | 15.6 | 2377 | 33 | 0.829 | 0.84 | 39 | 0.897 | 0.92 |
| 226 | 2.5 | 5.6 | 2384 | 25 | 0.806 | 0.65 | 28 | 0.894 | 0.66 |
| 242 | 7.7 | 3.5 | 2425 | 11 | 0.822 | 0.28 | 11 | 0.931 | 0.24 |
| 230 | 4.6 | 3.5 | 2446 | 12 | 0.734 | 0.33 | 9 | 0.834 | 0.22 |
| 234 | 10.6 | 12.5 | 2381 | 33 | 0.837 | 0.83 | 33 | 0.904 | 0.77 |
| 227 | 4.6 | 4.5 | 2328 | 15 | 0.808 | 0.40 | 14 | 0.903 | 0.33 |
| 316 | 2.8 | 4.6 | 2436 | 24 | 0.799 | 0.62 | 19 | 0.936 | 0.42 |
| 231 | 10.3 | 4.4 | 2255 | 25 | 0.809 | 0.69 | 26 | 0.935 | 0.62 |
| 318 | 20.7 | 14.8 | 2169 | 36 | 0.825 | 1.01 | 37 | 0.901 | 0.95 |
| 229 | 3.6 | 4.6 | 2172 | 10 | 0.806 | 0.29 | 10 | 0.908 | 0.25 |
| 314 | 3.6 | 3.4 | 2428 | 13 | 0.821 | 0.33 | 12 | 0.934 | 0.27 |
| 319 | 12.4 | 3.4 | 2411 | 9 | 0.800 | 0.23 | 10 | 0.898 | 0.23 |

Mean mutation rate from GATK calls: $0.59 \times 10^{-8}$ per bp, per generation
Mean mutation rate from *freebayes* calls: $0.53 \times 10^{-8}$ per bp, per generation

We found a strong association between paternal age and the number of *de novo* mutations inherited by offspring. For each additional year of paternal age at conception, offspring inherited an additional 1.5 *de novo* mutations (R^2^ = 0.79; Poisson regression, *p* = 2.9 × 10^-11^). Due to the structure of the pedigree among sampled individuals, we were able to unambiguously phase 139 of the 307 mutations (see Methods). Of these phased mutations, 76.3% (95% CI: [68.5, 82.6]) were determined to be of paternal origin. Figure 1 shows the count of phased mutations as a function of parental age, illustrating the large effect of paternal age on mutation rate (R^2^ = 0.78; Poisson regression, *p* = 7.5 × 10^-4^). In contrast to this relationship, we found no significant association between maternal age and either the number of mutations inherited or the estimated per-generation mutation rate (Fig. S2). Early studies of human trios also found no significant effect from maternal age (Kong et al. 2012; Francioli et al. 2015), though studies with much larger sample sizes have detected a small effect, potentially due to the accumulation of environmental damage in the maternal germline (Goldmann et al. 2016; Jónsson et al. 2017).

**Figure 1.**
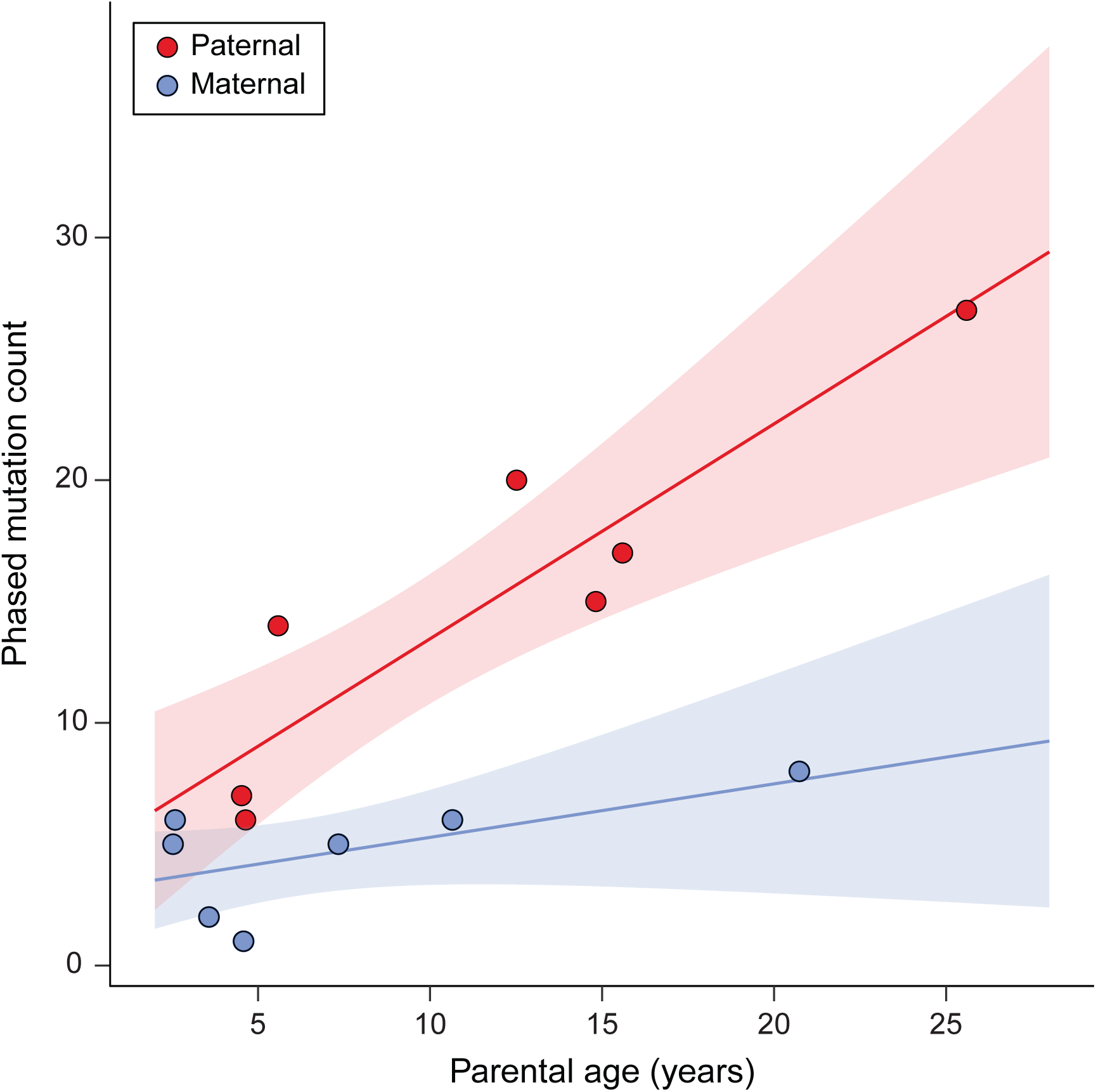
Phased mutation count and parental age. The number of phased mutations identified from seven rhesus macaque trios attributed to paternal (red) and maternal (blue) transmission. There is a strong linear relationship between the number of transmitted paternal mutations and the paternal age at conception (R^2^ = 0.78; Poisson regression *p* = 7.5 × 10^-4^). The number of maternally transmitted mutations was not significantly associated with the maternal age at conception in our data (R^2^ = 0.29; Poisson regression *p* = 0.11). Shaded areas show respective regression 95% CI. These seven trios represent cases in which mutations can be tracked through the following generation (Figure S1).

We found that C>T transitions at CpG sites accounted for 24% of all observed point mutations (similar to the 17% and 22-24% reported in humans and chimpanzees, respectively; Kong et al. 2012; Venn et al. 2014; Besenbacher et al. 2019). CpG sites are more prone to mutation than other sites due to spontaneous deamination (Bird 1980). We estimated the mutation rate at CpG sites to be 1.43 × 10^-7^ per bp per generation, an order of magnitude higher than the rest of the genome. The number of mutations at CpG sites was also significantly associated with paternal age (R^2^ = 0.20, Poisson regression, *p* = 0.015; Fig. S3).

The mutation spectrum in rhesus macaques is similar to that found in other primates, except for a slight excess of C>T transitions (Fig. 2). C>T transitions accounted for 54% of all observed *de novo* mutations, significantly higher than the 42% found in human (χ^2^ test, *p* = 1.5 × 10^-4^; Jónsson et al. 2017). This excess in C>T transitions results in a higher transition-to-transversion ratio in macaque (2.84) than observed in humans (2.22). This excess also contributes to a higher mutation rate toward weak base pairs (A:T) versus strong base pairs (G:C) when compared to humans (macaque: 3.51, human: 2.11).

**Figure 2.**
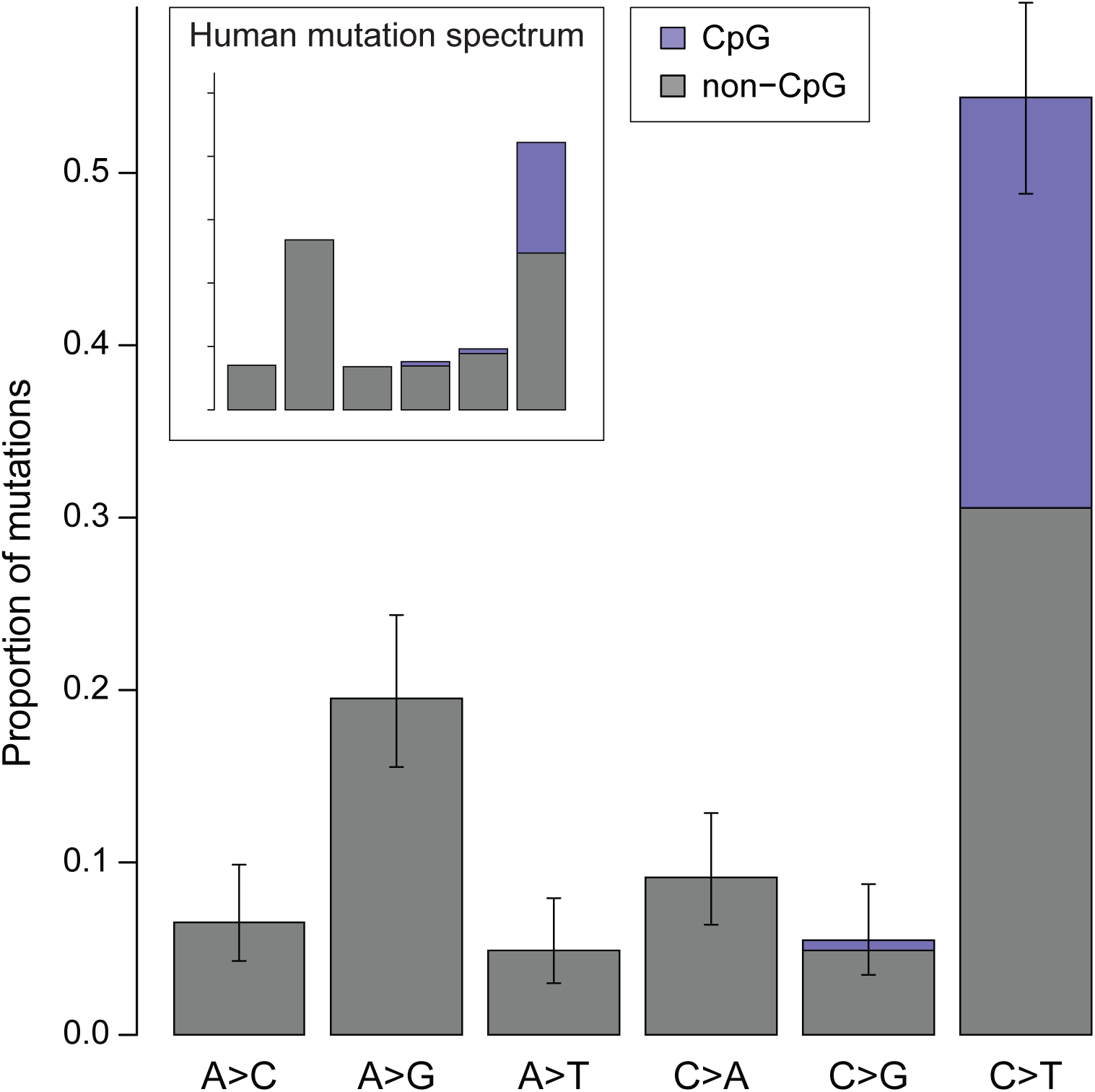
Mutation spectrum in rhesus macaque. The frequency of each type of mutation from among the 307 identified. Error bars show binomial proportion 95% CI (Wilson score interval) for totals at each type. Mutations at CpG sites accounted for 24% of all mutations and represent 43% of all strong-to-weak transitions. Mutation categories represent their reverse complement as well.

### A lower per-generation mutation rate in the rhesus macaque

Our overall estimate of the per-generation mutation rate in rhesus monkeys (0.58 × 10^-8^ per bp per generation) is lower than direct estimates from other primates, but both the average age of parents and the average age at puberty differs among species (Table 2). The average paternal age at conception explains most of the variation among studies in direct estimates of the human mutation rate (Kong et al. 2012; Rahbari et al. 2016; Jónsson et al. 2017). A meaningful comparison of mutation rates between species must therefore consider variation in the paternal age at conception among sampled individuals. Figure 3 shows the rate of mutation accumulation with paternal age at conception in macaque trios compared to the rate observed in humans. To compare the mutation rate estimate from rhesus macaques to other species while also accounting for different average ages at puberty between species, we considered a model of reproductive longevity (Thomas et al. 2018).

**Figure 3.**
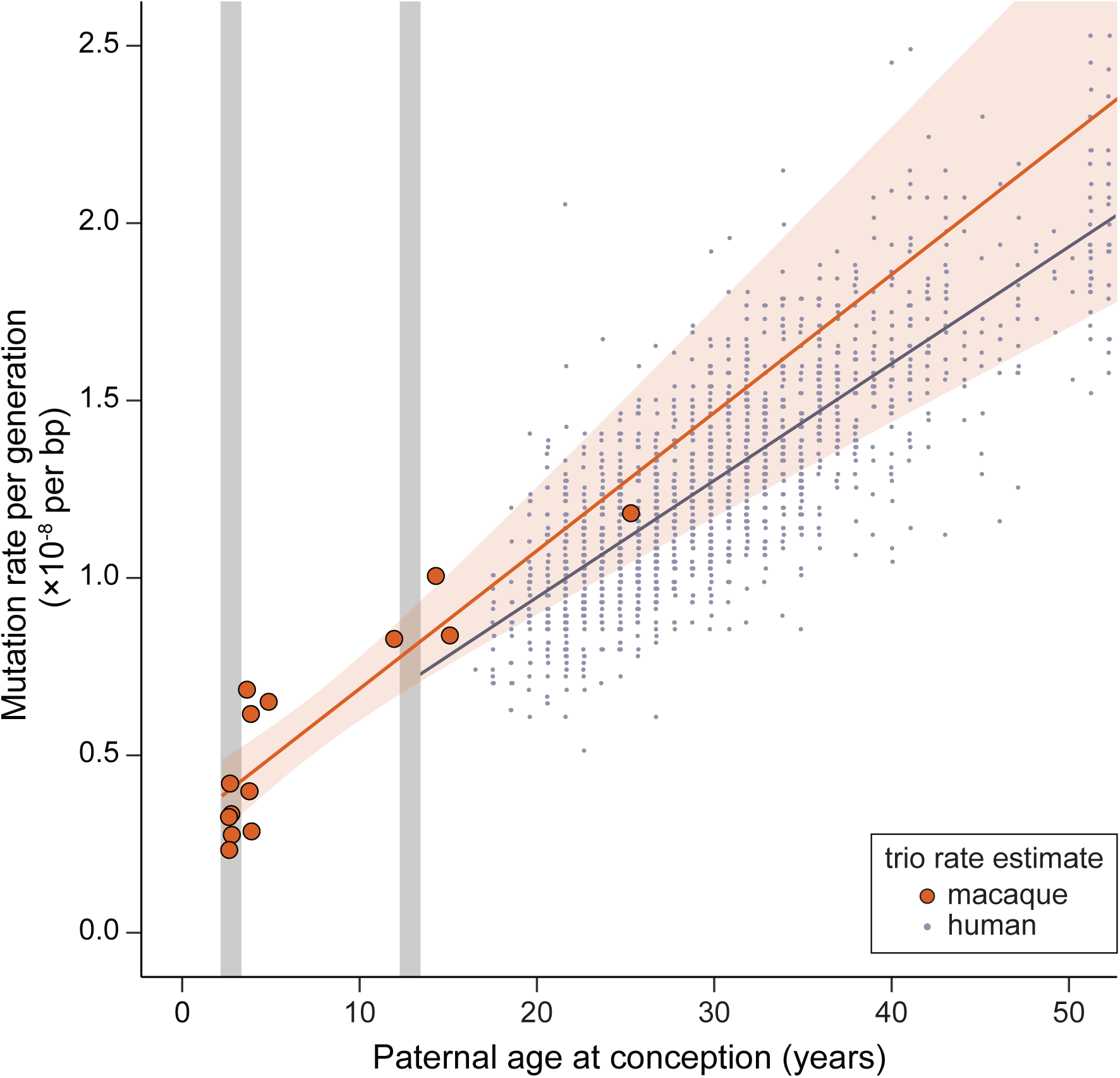
Similar rates of mutation accumulation post-puberty in human and rhesus macaque. Mutation rate accumulation with paternal age estimated from trios in macaques (orange) and humans (black, data from Jónsson et al. (2017)). Approximate age at male puberty in the macaque (3.5 y) and human (13.5 y) are shown in grey. Human trios with paternal age up to 50 are shown here, but the human regression line is from the full dataset. The rate at which the mutation rate increases with paternal age is slightly higher in the macaque (4.3 × 10^-10^ per bp, per year; Poisson regression) than in human (3.4 × 10^-10^ per bp, per year). The intercept with puberty is much lower in macaque (3.9 × 10^-9^ per bp) than in human (7.1 × 10^-9^ per bp).

**Table 2.**
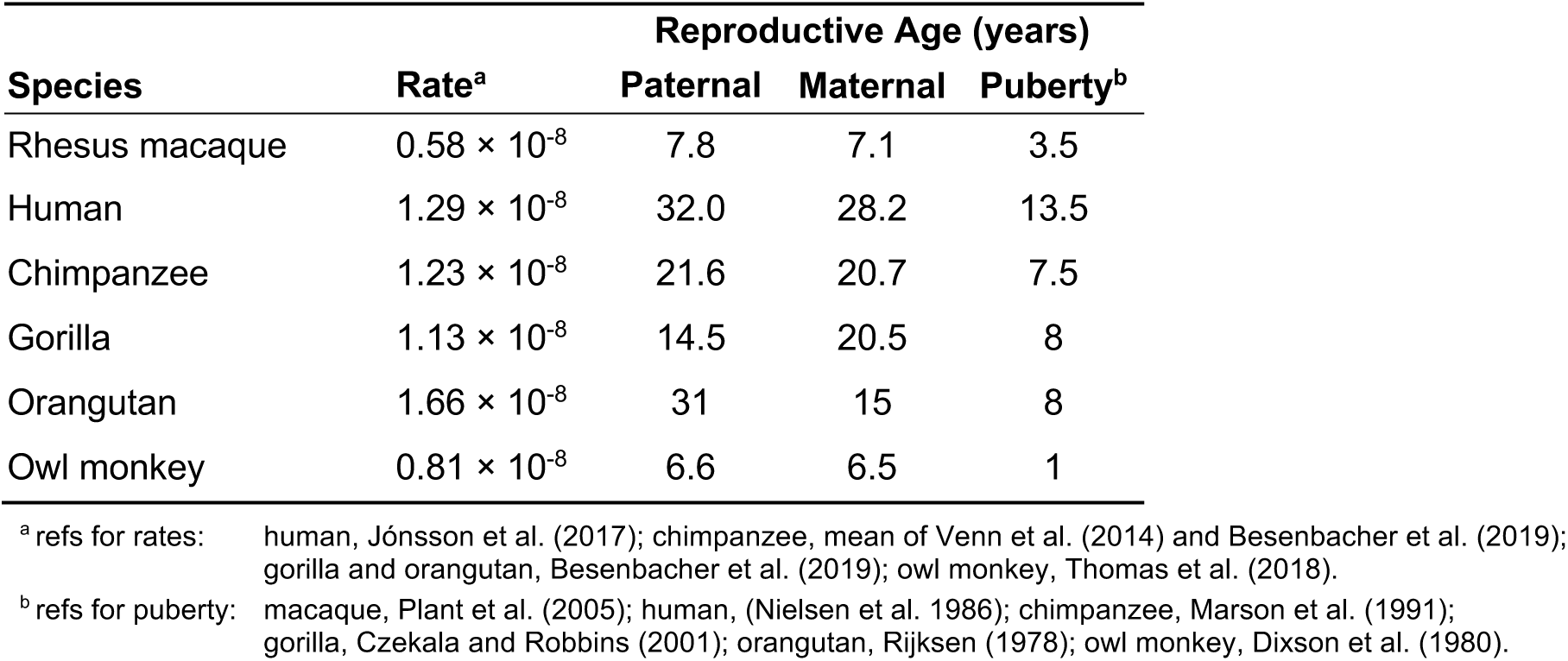
Per-generation mutation rate and reproductive age in primate studies

| **Species** | **Rate<sup>a</sup>** | **Reproductive Age (years)** |  |  |
| --- | --- | --- | --- | --- |
|  |  | **Paternal** | **Maternal** | **Puberty<sup>b</sup>** |
| Rhesus macaque | $0.58 \times 10^{-8}$ | 7.8 | 7.1 | 3.5 |
| Human | $1.29 \times 10^{-8}$ | 32.0 | 28.2 | 13.5 |
| Chimpanzee | $1.23 \times 10^{-8}$ | 21.6 | 20.7 | 7.5 |
| Gorilla | $1.13 \times 10^{-8}$ | 14.5 | 20.5 | 8 |
| Orangutan | $1.66 \times 10^{-8}$ | 31 | 15 | 8 |
| Owl monkey | $0.81 \times 10^{-8}$ | 6.6 | 6.5 | 1 |

<sup>a</sup> refs for rates: human, Jónsson et al. (2017); chimpanzee, mean of Venn et al. (2014) and Besenbacher et al. (2019); gorilla and orangutan, Besenbacher et al. (2019); owl monkey, Thomas et al. (2018).
<sup>b</sup> refs for puberty: macaque, Plant et al. (2005); human, (Nielsen et al. 1986); chimpanzee, Marson et al. (1991); gorilla, Czekala and Robbins (2001); orangutan, Rijksen (1978); owl monkey, Dixson et al. (1980).

Reproductive longevity is defined as the amount of time a parent is in a reproductive state prior to offspring conception: here, we use the paternal age at conception minus the age at sexual maturity (i.e. the age at puberty). In this model, the mutation rate can evolve between species if the rate of replication-dependent errors changes, if the rate of germline cell-division post-puberty changes, or if there are different numbers of mutations that accumulate prior to puberty (Thomas and Hahn 2014; Thomas et al. 2018). Though key parameters of spermatogenesis surely differ between species, a simple model that considers only the difference between the beginning of spermatogenesis and the average age at conception makes a compelling null model for understanding mutation rate variation.

We performed a Poisson regression of mutation count on parental age to model the rate of accumulation with reproductive longevity, and compared it to a regression from a large human dataset (Jónsson et al. 2017). Because the regression was not significant for maternal age in the macaque data, we excluded maternal age as a variable for the subsequent comparisons. If we assume both that the number of mutations before puberty and the rate of accumulation of mutations after puberty are the same in humans as in macaques, our model with reproductive longevity overestimates the expected number of mutations per generation (approx. 53 vs. 36, using 7.8 years as the average paternal age in macaques).

Much of the difference in the per-generation mutation rate between human and macaque can be attributed to the number of mutations predicted in the germline pre-puberty: there are about half as many in macaques as in humans (25.4, 95% CI: [21.0, 29.7] in macaque and 44.1 [43.0, 45.2] in human, based on regression predictions at species’ respective timing of puberty). The rate at which mutations increase with paternal age after puberty was not significantly different between macaque (4.3 × 10^-10^ per bp, per year, 95% CI [3.0, 5.5]) and human (3.4 × 10^-10^ per bp, per year, [3.3, 3.5]; unequal variances *t*-test, *p* = 0.17). The non-significant, but slightly higher effect of paternal age in macaques corresponds to only ∼2 more mutations over the average lifespan of a macaque.

The evolutionary changes leading to a lower number of mutations before puberty appear to have occurred along the branch leading to macaques, as owl monkeys (*Aotus nancymaae*) show a similar number of mutations pre-puberty as humans (Thomas et al. 2018). The smaller number of mutations in macaques may be due to a decreased number of germline cell divisions or a decreased rate of mutation per cell division before puberty. Whichever the case, the data here suggest that the mutation rate per cell division post-puberty has had little effect on the overall difference in the per-generation rate between species.

We further used the regression model to estimate a per-year mutation rate for the macaque. Such values can be directly compared to substitution rates from phylogenetic studies. Instead of dividing the per-generation rate by the average parental age from our sample, we calculated the per-year rate as a property of the species by predicting the mutation rate at the median age of reproduction. For a paternal age of 11 years in macaques (Xue et al. 2016), our regression model predicts a mutation rate of 0.71 × 10^-8^ per bp per generation for macaques, corresponding to a rate of 0.65 × 10^-9^ per bp per year. This per-year rate is higher than the × 10^-9^ per bp per year observed in humans (Jónsson et al. 2017), consistent with reports of a lower human per-year mutation rate (Scally and Durbin 2012; Ségurel et al. 2014; Besenbacher et al. 2019).

### Sociability in male rhesus monkeys shows no connection with sire age

Sociability is a consistent personality dimension in humans that has also been identified in rhesus monkeys (Capitanio 1999). Low social ability in infant rhesus monkeys has been shown to predict poor adult social function (Sclafani et al. 2016), consistent with deficits in childhood social interaction and communication as risk factors for the development of autism spectrum disorder in humans (Ozonoff et al. 2010; Jones et al. 2014). We examined sociability across a sample of 203 male monkeys studied in adulthood to determine whether paternal age at conception was a significant contributor to low social function. These monkeys came from the same colony as those used for sequencing, but none of the individuals were the same.

In addition to sociability, we measured the frequency of eight behaviors associated with general social functioning, stratified by sex (Table S1). We performed principal component analysis on these variables to reduce dimensionality and to extract a useful general score of social functioning from these behaviors. The first two principal components (PC) explain > 94% of the variance in these observations. Offspring social behavior PC1 explains the tendency for behaviors to be directed towards females versus males, while offspring social behavior PC2 explains overall contact and proximity with both sexes. Social behavior PC2 was significantly correlated with observer ratings of the sociability personality measure (Pearson’s *r* = 0.37, *p* < 5 × 10^-9^).

We found no evidence for a relationship between paternal age and any measure of lowered social function (Fig. 4). Rather than a negative effect on sociability, there was a weak positive trend suggested between sociability and parental age at conception (sire age: *r* = 0.07, *p* = 0.21; dam age: *r* = 0.02, *p* = 0.09). Because there is a positive correlation between sire rank and sire age, we also calculated pairwise partial correlations between sire age and all measures of social functioning while attempting to control for sire rank. None of these correlations were significant (Table S2).

**Figure 4.**
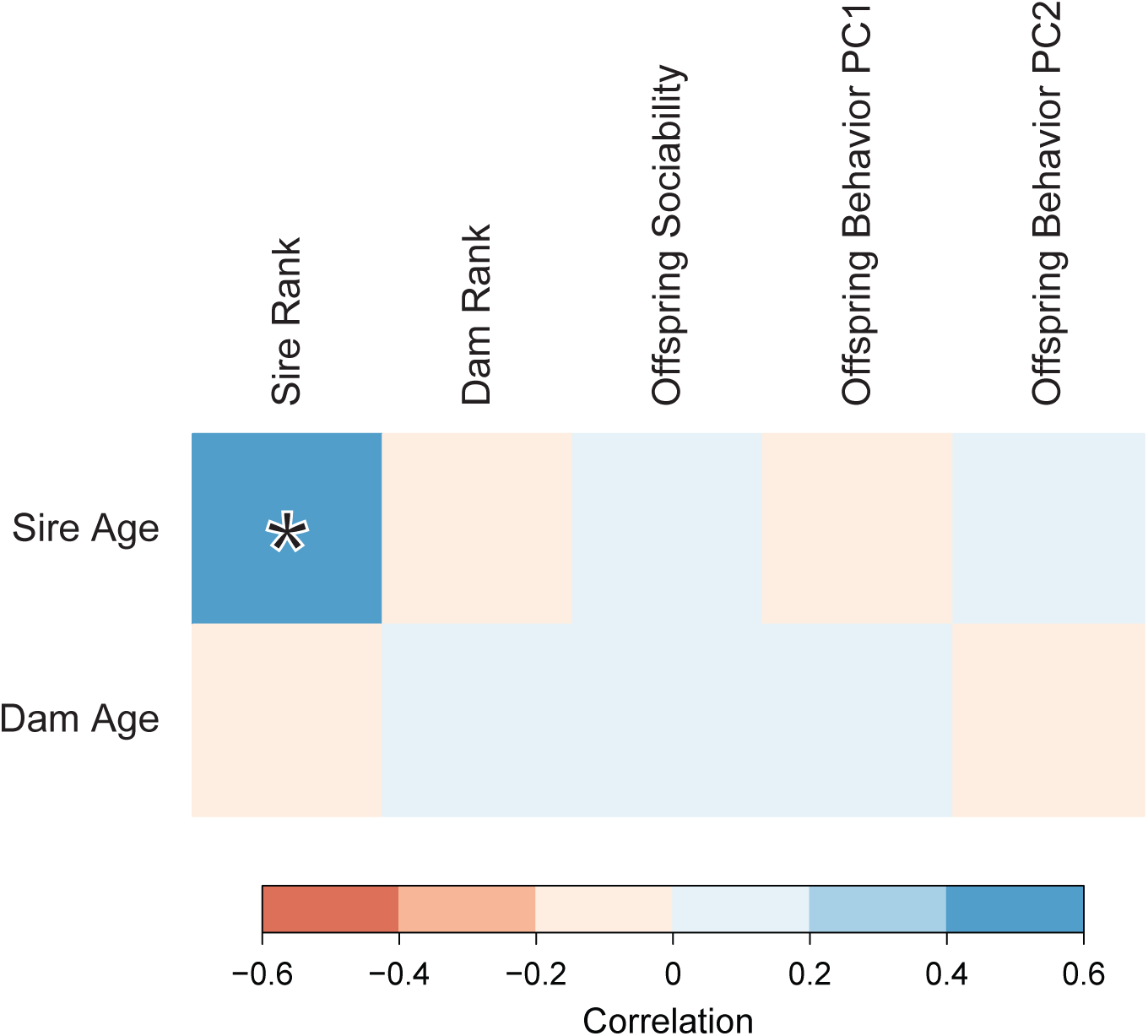
Correlations between parental age and behavioral traits in male rhesus monkeys. Boxes are shaded by the intensity of correlation in pairwise comparisons. Legend shows range of Pearson’s correlation coefficient for each color. Significant correlation (*p* < 0.05) highlighted with an asterisk.

## Discussion

Both the rate and the spectrum of mutations are intimately linked with life history (Walter et al. 2004; Goldmann et al. 2016; Rahbari et al. 2016), complicating comparisons across studies that report point estimates. We discovered a significant paternal age effect on mutation rate in rhesus macaques, consistent with findings from other direct estimates of the mutation rate in primates. The overall per-generation mutation rate in the macaque is substantially lower than has been found in humans and other great apes, but similar to the rate in owl monkeys. Our analysis indicates that this lower value compared to the apes is largely due to a younger age at reproduction and a lower number of mutations before puberty, with little effect from differences in the rate at which mutations accumulate with paternal age after puberty. While the effect of maternal age was positive, it was not significant. Nevertheless, the ratio of male-to-female mutations, *α* ≈ 3, closely matches the value described in human studies (Jónsson et al. 2017; Gao et al. 2019). The stability of *α* with paternal age is expected if maternal and paternal mutations are both increasing linearly with age. The similarity in *α*-values, coupled with the small but significant effect of maternal age on mutation rate in humans, suggests that a maternal age effect might be detected in macaques with larger sample sizes.

In contrast to the per-generation rate, our estimate of the per-year mutation rate in macaques is 1.5 times higher than the estimate in humans. This factor is similar to recent reports that the per-year mutation rate is roughly 1.4-1.5 times higher in chimpanzees, gorillas, and orangutan relative to humans (Besenbacher et al. 2019). Estimates from phylogenetic studies, however, indicate only a 30% higher per-year substitution rate in Old World monkeys (including macaques) compared to humans (Kim et al. 2006; Elango et al. 2009). This discrepancy is consistent with a recent, and perhaps ongoing, slowdown in the mutation rate on the human lineage (Goodman 1985; Li and Tanimura 1987; Yi 2013). That is, if a lower per-year mutation rate evolved sometime after the human-chimpanzee divergence, then a comparison of substitution rates that uses the entire divergence time—as in the Old World monkey to human comparison—will underestimate the degree to which the rate has decreased. The estimated 1.5-fold higher per-year mutation rate in chimpanzee relative to human also supports the recent evolution of a lower per-year mutation rate specific to the human lineage, though it is not consistent with the much smaller difference in substitution rate between these two species (Yi 2013).

Despite accounting for the number of mutations transmitted with paternal age, a model that adjusts for reproductive longevity (Thomas et al. 2018) does not account for all differences in mutation rate between macaques and humans. The biggest difference between these species appears to be the number of mutations present at puberty, before active spermatogenesis begins. It is not clear, however, what changes have occurred before puberty to lower the mutation rate in macaques. Though our data suggest that the mutation rate per-cell division post-puberty is the same between species, it is possible that there are differences between species in the error-prone divisions of early embryogenesis (Huang et al. 2014; Rahbari et al. 2016; Ju et al. 2017). Under such a model, the decreased number of mutations before puberty in the macaque may be explained by a lower number of postzygotic mutations, a process that is not modeled well by mutation rates during spermatogenesis. In any case, the evolution of life-history appears to have played a large role in shaping differences in the per-generation mutation rate between human and macaque.

With the large effect of paternal age on mutation rates within species, differences in key parameters of spermatogenesis—including timing of cell division, cell cycle length, and efficiency—have been hypothesized to explain variation in mutation and substitution rates between species (Wilson Sayres et al. 2011; Thomas and Hahn 2014; Amster and Sella 2016; Moorjani et al. 2016; Scally 2016). The seminiferous epithelial cycling time is one such parameter of particular interest because it makes straightforward predictions about mutation rate variation between species. This time describes the period between successive waves of spermatogenesis in the testis, and is known to be 34% shorter in the macaque than in humans (de Rooij et al. 1986). All things being equal, the shorter time between cycles suggests that male macaques should accumulate mutations post-puberty at a higher rate than male humans. However, our results reveal little difference between macaques and humans in how mutation accumulation scales with paternal age. The absence of a proportionate effect on mutation rate brings into question the long-held assumption that spermatogonial stem cells (SSCs) are all actively dividing (Drost and Lee 1995). If division by only a fraction of SSCs are needed to replenish the seminiferous epithelium each cycle, a shorter cycling time would not require a commensurate increase in the number of stem cell divisions. Cell proliferation experiments in the macaque suggest that the fraction of SSCs actively dividing varies under endocrine control (Simorangkir et al. 2009; Plant 2010). Relaxing the assumption that spermatogonial stem cells are all actively dividing may also help to explain the reported disconnect between the male-to-female ratio of germline divisions and the ratio of X-to-autosome substitutions (Wilson Sayres and Makova 2011; Ségurel et al. 2014; Gao et al. 2016; Scally 2016).

While our finding of a linearly increasing number of mutations with paternal age in the macaque is consistent with a replication-dependent model of mutagenesis, a role for damage-dependent mutagenesis cannot be ruled out. For instance, a model of mutagenesis that relies solely on environmental damage could explain why differences in the seminiferous epithelial cycling time between species have no effect on the rate of mutation accumulation with paternal age. However, such a model is difficult to reconcile with the differences in substitution rates seen across primates. A damage-dependent model also need not be independent of the rate of cell division, as both the male mutation bias and paternal age effect can be explained if the accumulation of spontaneous mutations relies on the rate of cell division (Gao et al. 2016; Seplyarskiy et al. 2019). In such a model, the repair of DNA lesions is limited by the time between replications: rapidly dividing cells such as spermatocytes accumulate a greater number of mutations because there is less time for lesions to be repaired. The replication process instead ensures that such lesions appear as mutations in future generations. The presence of a paternal age effect on mutations at CpG sites (e.g. Fig. S3), despite their ostensibly replication-independent origin, is better explained by a model of unrepaired damage before replication.

Previous studies have found that both the number of *de novo* mutations and the risk of neurodevelopmental disorders increase with paternal age in humans (Kong et al. 2012). We find no link between paternal age and negative social behavioral outcomes in offspring, despite an increasing number of mutations with paternal age in the rhesus macaque. We must acknowledge that social behavior is a complex human construct that our assay is unlikely to fully capture. Furthermore, differences in sociability are only a single component in the complex syndromes that constitute neurodevelopmental disorder. Nevertheless, our findings are consistent with hypotheses that the increased risk of neurodevelopmental disorder with paternal age in humans is not primarily driven by *de novo* mutations (Hultman et al. 2011; Gratten et al. 2016; Janecka et al. 2017). While these hypotheses do not exclude a role for inherited genetic factors in the development of such disorders, they posit no direct role for the elevated *de novo* mutation rate in the higher risk of disorders observed in offspring of older fathers.

## Methods

### Sequencing

Genomic DNA isolated from blood samples (buffy coats) from 32 Indian-origin rhesus macaques from the California National Primate Research Center (Univ. of California at Davis) was used to perform whole genome sequencing. We generated standard PCR-free Illumina paired-end sequencing libraries using KAPA Hyper PCR-free library reagents (KK8505, KAPA Biosystems) in Beckman robotic workstations (Biomek FX and FXp models). We sheared total genomic DNA (500 ng) into fragments of approximately 200-600 bp using the Covaris E220 system (96-well format) followed by purification of the fragmented DNA using AMPure XP beads. We next employed a double size selection step with different ratios of AMPure XP beads to select a narrow size band of sheared DNA molecules for library preparation. DNA end-repair and 3′-adenylation were performed in the same reaction followed by ligation of the barcoded adaptors to create PCR-Free libraries. We ran each library on the Fragment Analyzer (Advanced Analytical Technologies, Ames, Iowa) to assess library size and presence of remaining adaptor dimers. This was followed by qPCR assay using KAPA Library Quantification Kit and their SYBR FAST qPCR Master Mix to estimate the size and quantification. These WGS libraries were sequenced on an Illumina HiSeq-X instrument to generate 150 bp paired-end reads. All flow cell data (BCL files) were converted to barcoded FASTQ files.

### Mapping and variant calling

BWA-MEM version 0.7.12-r1039 (Li 2013) was used to align the Illumina sequencing reads to the rhesus macaque reference assembly Mmul_8.0.1 (GenBank accession GCA_000772875.3) and to generate BAM files for each of the 32 individuals. Picard MarkDuplicates version 1.105 (Broad Institute 2019) was used to identify and mark duplicate reads. Single nucleotide variants (SNVs) and small indels (up to 60 bp) were called using GATK version 3.6 (Van der Auwera et al. 2013) following best practices. We used HaplotypeCaller to generate gVCFs for each sample, and performed joint genotype calling on all samples using GenotypeGVCFs, generating a VCF file. GATK hard filters (SNPs: “QD < 2.0 || FS > 60.0 || MQ < 40.0 || MQRankSum < −12.5 || ReadPosRankSum < −8.0”; Indels: “QD < 2.0 || FS > 200.0 || ReadPosRankSum < −20.0”) were applied and we removed calls that failed.

We also applied an alternative pipeline, *freebayes* version v0.9.21-19-gc003c1e (Garrison and Marth 2012), to get an independent estimate of *de novo* mutations. We ran *freebayes* on the BAM files and applied the recommended filters to the resulting calls: (“QUAL > 1 & QUAL / AO > 10 & SAF > 0 & SAR > 0 & RPR > 1 & RPL > 1”).

### Filtering candidate mutations

Our initial list of candidates from GATK variant calls included 300,065 Mendelian violations (MVs) among the 14 trios. The initial list from *freebayes* calls included 23,240 MVs. In the next steps, we progressively applied filters that reduced both the number of candidates and the total number of callable sites to improve the accuracy of identifying actual *de novo* mutations. To be specific, we:

- Removed candidate sites that had fewer than 20 reads or more than 60 reads. Sites with too few reads have a higher chance of being undersampled, while sites with too many reads may represent problematic repetitive regions (Li 2014).
- Removed candidate sites that were not homozygous reference in both parents, or appeared as a variant in an unrelated trio. This latter step reduces the chances that a child inherited a segregating variant that was miscalled as homozygous reference in a parent.
- Restricted candidate sites to those with high genotype quality (GQ > 70) in both parents and offspring (see Fig. S6).
- Removed heterozygotes in the offspring that did not have at least 1 alternate read on both the forward and reverse strand (i.e. ADF > 0 and ADR > 0). We also removed candidate sites where homozygote calls in the parent had more than 1 alternate read on either strand (AD < 2). These alternate allelic depths were evaluated before genotype calling to minimize genotyping errors from local realignment (Karczewski et al. 2019).
- Removed heterozygotes in the offspring that were called with an allelic depth of less than 35% alternate reads (Fig. S7).

The same filters were applied to MVs from both variant calling pipelines, except for GQ which is not calculated by default in *freebayes*. To evaluate the sensitivity of our mutation calls to the GQ filter, we re-estimated the mutation rate (accounting for callability, see below) at several filter limits (Fig. S5). After applying the above filters to the set of MVs from both pipelines, we found: 269 overlapping candidates, 44 unique to *freebayes*, and 38 unique to GATK. Our subsequent analyses use the set of mutations from the GATK calls, but estimated mutation rates are similar between calls from the two pipelines after accounting for differences in callability (Table 1).

### Estimating the fraction of callable sites

To calculate a mutation rate while considering differences in coverage and filtering, we adapted the strategy from Besenbacher et al. (2019). Raw counts of *de novo* mutations were converted into a mutation rate by dividing by the total number of callable sites. Estimates of site callability, the probability that a true *de novo* mutation would be correctly called as such at a given site *x*, are factored into estimates of the mutation rate by using the following equation:

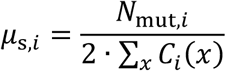

where *μ*_s,*i*_ is the per-site per-generation mutation rate for trio *i*, *N*_mut,*i*_ is the number of *de novo* mutations identified in trio *i*, and *C_i_*(*x*) is the callability of site *x* in that trio. We take *x* to be a site from the set of all haploid sites with depth between 20 and 60. This strategy assumes that the ability to correctly call each individual in the trio is independent, allowing us to estimate *C_i_*(*x*) as:

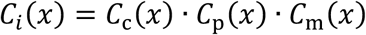

where *C*_c_, *C*_p_, and *C*_m_ are the probability of calling the child, father, and mother correctly in trio *i*. We estimate each of these by considering the proportion of sites that pass our set of filters in a set of high-confidence calls from each trio. For heterozygous calls in the child, we estimate:

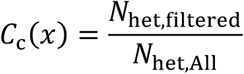

taking *N*_het,All_ to be the number of variants where one parent is homozygous for the reference allele and one parent is homozygous for the alternate allele with high confidence. *N*_het,filtered_ is the number of heterozygote calls in the child remaining after applying all filters (including ADF > 0, ADR > 0, and alternate allele depth > 35%).

Similarly, we estimate parental callability as (in the case of the father):

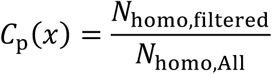

where *N*_homo,All_ is the number of variants where both parents are homozygous for the reference allele with high confidence and *N*_homo,filtered_ is the number of homozygous calls in the child that pass all filters (including AD < 2). To best match these homozygous parental sites with candidate *de novo* sites, sampled sites were restricted to those where the variant was present no more than once across all individuals (i.e. allelic count, AC < 2).

We estimated callability for each individual, *C*_c_, *C*_p_, and *C*_m_, from a random sample of 250,000 sites across the genome that matched the respective criteria for each trio. The *de novo* callability, *C_i_*(*x*), calculated for each trio from this strategy is listed in Table 1.

### Phasing mutations

We traced the parent of origin for *de novo* mutations that were transmitted to the third generation. This was accomplished by tracking their inheritance on haplotype blocks that we assembled from phase-informative sites. These informative sites were biallelic and had genotypes that were different between grandparents, heterozygous in the next generation, and not heterozygous in both the third-generation proband and the other parent. These phase-informative sites could be traced unambiguously to one of the grandparents. Sites were assembled into haplotype blocks under the assumption that multiple recombination events in a single meiosis were unlikely to occur within a 1-Mb interval (Rogers et al. 2006; Huang et al. 2009; Smeds et al. 2016; Xue et al. 2016).

### Mutation rates with parental age

We estimated the effect of parental age on the mutation rate with a Poisson regression, modeling the number of mutations for trio *i* as:

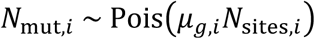

where *μ*_g*,i*_ is the per-generation mutation rate in trio *i* and *N*_sites,*i*_ *=* 2 · ∑_6_ *C*_*i*_(*x*), the diploid callable genome size for trio *i*. We used an identity link in the Poisson regression, which fit better than the canonical link (lower AIC value), with:

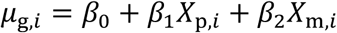

where *X*_p_ and *X*_m_ are the paternal and maternal ages for trio *i,* and respective regression coefficients *β*. To adjust for differences in observable genome size, we generated two new variables:

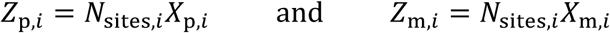

and fit the Poisson regression model on *N*_sites,*i,*_ *Z*_p,*i,*_ and *Z*_m,*i*_ with no intercept. That is:

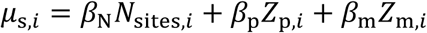

where *μ*_s*,i*_ is the per-site per-generation mutation rate in trio *i*.

In this regression model, the coefficient for *N*_sites,*i*_ represents the per-site version of the original intercept *β*_0_, and the coefficients for *Z*_p,*i*_ and *Z*_m,*i*_ are the effects of the respective parental age on the per-site per-generation mutation rate.

We compared our Poisson regression model for the mutation rate in rhesus macaque to a model of the mutation rate in human. Since maternal age was not significantly correlated with mutations in macaque, we omitted the variable in the comparison between species. This yielded the following regression coefficients: *β*_N_ = 2.43 × 10^-9^, 95% CI [1.47, 3.38], *β*_P_ = 4.27 × 10^-10^ [3.01, 5.53] for macaque and *β*_N_ = 2.56 × 10^-9^ [2.26, 2.86], *β*_P_ = 3.37 × 10^-10^ [3.28, 3.47] for human. To test whether coefficients in this regression were significantly different, we used an unequal variances *t*-test. This test statistic was equal to the absolute difference between coefficients divided by the pooled standard error.

We predicted the number of mutations at different paternal ages using the above regression model and a diploid genome size (i.e. *N*_sites,*i*_) approximated by twice the UCSC golden path length for each species (genome.ucsc.edu). If the same number of mutations accumulate before puberty in rhesus macaque as in human, and the rate of accumulation after puberty is the same, our sample of macaques should have the same number of mutations as an average 17.8-year-old human. This is based on the mean paternal age of 7.8 years among macaques in our dataset and an age of 3.5 years for male puberty in macaques (Plant et al. 2005), resulting in 4.3 years of post-puberty mutation accumulation. Together with the 13.5 years to reach male puberty in humans, the corresponding age for the human model becomes 17.8 years (=13.5 + 4.3).

### Collecting sociability data from captive rhesus monkeys

We observed 203 male rhesus monkeys at the California National Primate Research Center, in cohorts of 5 to 8 animals (mean age = 6.9 years, range = 4.0 to 19.2 years), unobtrusively in their half-acre outdoor enclosures across four summers. Observations were conducted over the course of eight days within a two-week period and consisted of two 10-minute sessions per day. Using focal animal sampling (Altmann 1974), behavioral observers recorded the frequencies of the following behaviors directed at other adult animals: approach (locomotion to within arm’s reach), proximity (being within arm’s reach for at least three seconds), contact (physical, non-aggressive contact between animals), and grooming (picking with fingers and/or licking another animal’s hair). Following completion of behavioral observations on each cohort, the observer rated each animal using a 7-point Likert-type scale on three trait adjectives. Previous work (Capitanio and Widaman 2005) had demonstrated these ratings form a scale that reflected personality characteristics as follows: Sociability—affiliative, agreeable, sociable, appears to like the company of others and seeks out social contact with other animals; Warm—seeks or elicits bodily closeness, touching, grooming; and Solitary—subject prefers to spend considerable time alone, avoids or does not often seek contact with other animals. Observers were trained to show greater than 85% agreement on coding behavior, and Cronbach’s alpha (a measure of scale reliability) was greater than 0.92 for all samples.

## Data Access

A table with details on all identified *de novo* mutations is available as Supplemental Table 3. All sequencing data generated in this study has been submitted to the NCBI Sequence Read Archive under accession number: <Uploaded to SRA, will be released upon publication>

## Acknowledgments

This work was funded by the Precision Health Initiative of Indiana University. Behavioral assessments were made possible by funding from CNPRC base grant P51 OD011107 and NIH grants R37 AG033590 and R24 OD010962. We would like to acknowledge the production staff of the Human Genome Sequencing Center and its Director, Richard Gibbs. Three reviewers gave valuable feedback that greatly improved the manuscript.

---

**Figure S1.**
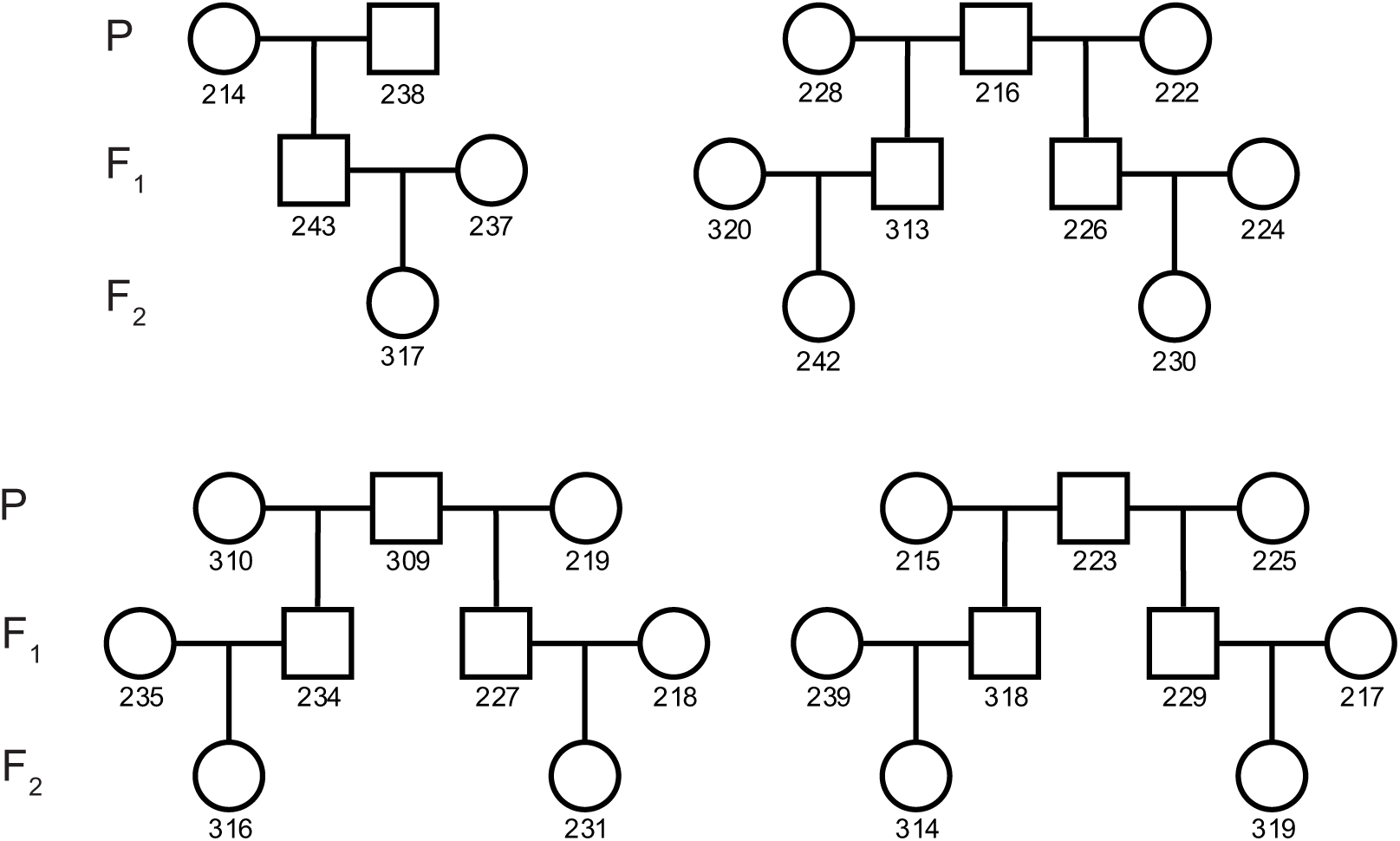
Pedigree structures of sequenced macaques. The 32 individuals sampled in this study were each part of the three-generation families depicted above.

**Figure S2.**
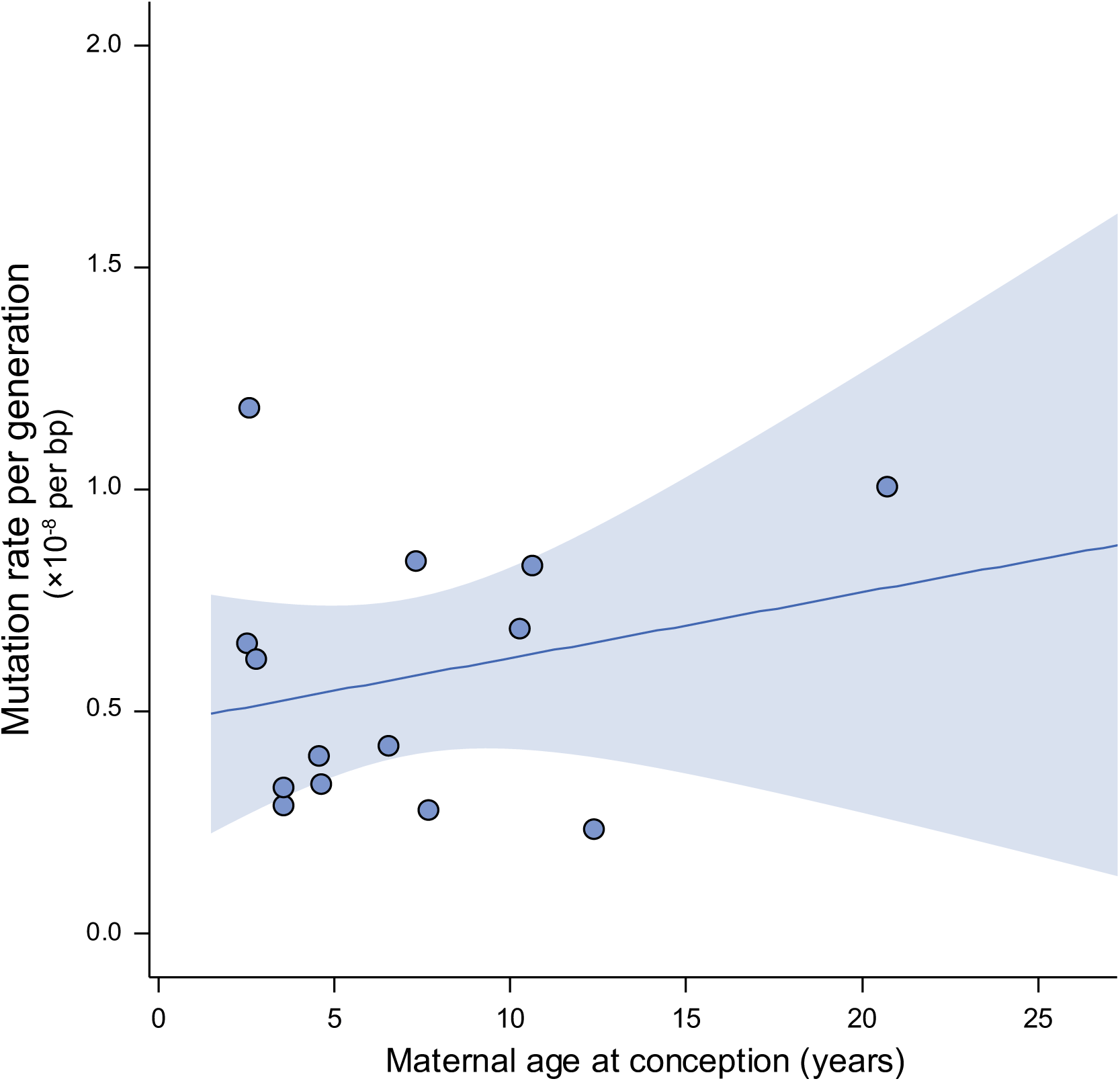
Effect of maternal age on mutation rate. Estimates of the overall mutation rate from 14 independent rhesus macaque trios with respect to the maternal age at conception. No significant relationship exists between overall mutation rate and maternal age at conception (R^2^ = 0.02, *p* = 0.39; shaded area shows 95% CI).

**Figure S3.**
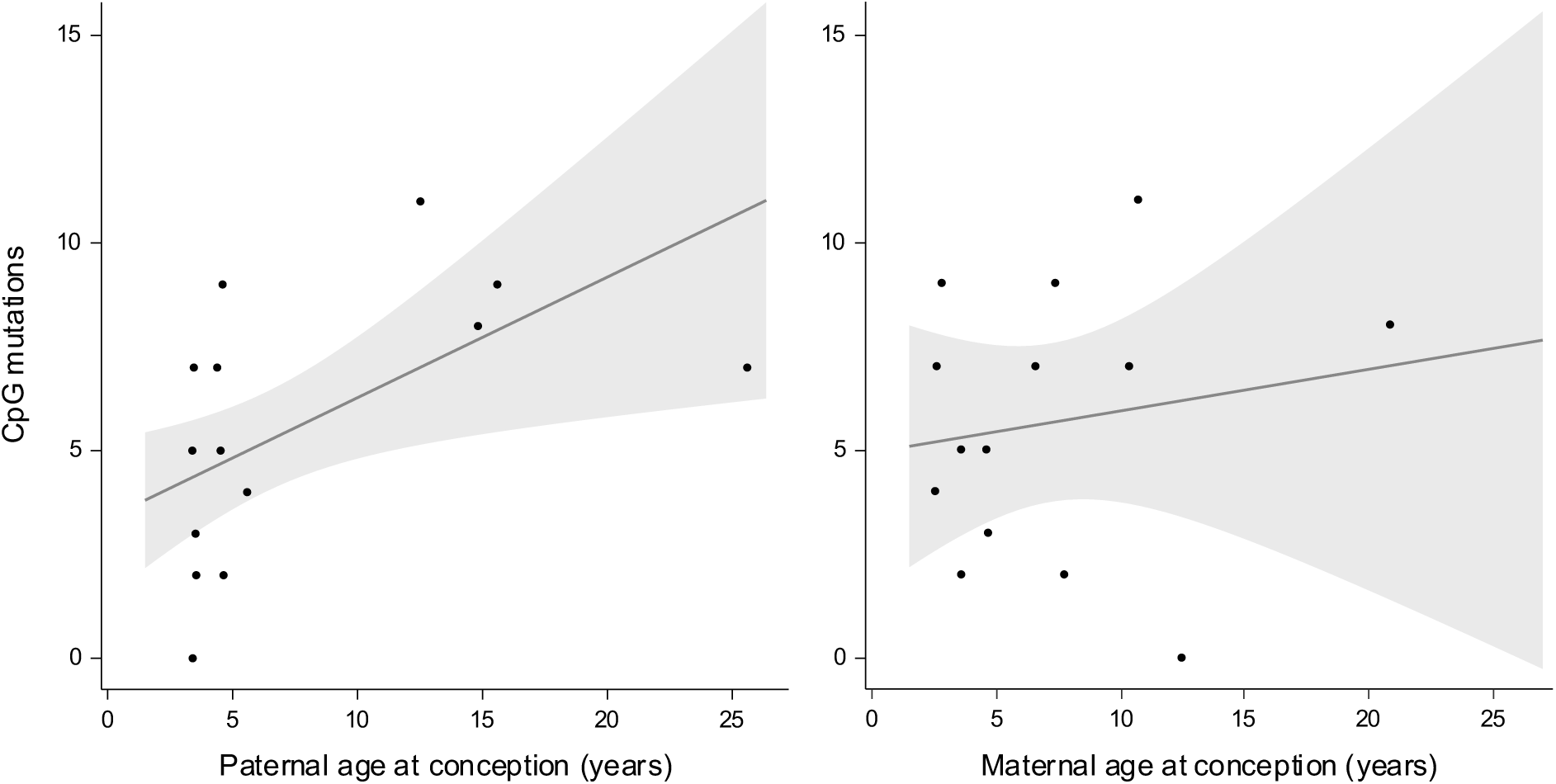
Mutations at CpG sites and parental age. Mutations at CpG sites with respect to the paternal (left) and maternal (right) age in each of the 14 trios. Shaded area shows regression 95% CI. As with the overall mutation rate, there is a significant relationship between the CpG mutation rate and paternal age (R^2^ = 0.27; Poisson regression, *p* = 0.015) while we find no evidence for a relationship with maternal age (R^2^ = 0.03; Poisson regression, *p* = 0.94).

**Figure S4.**
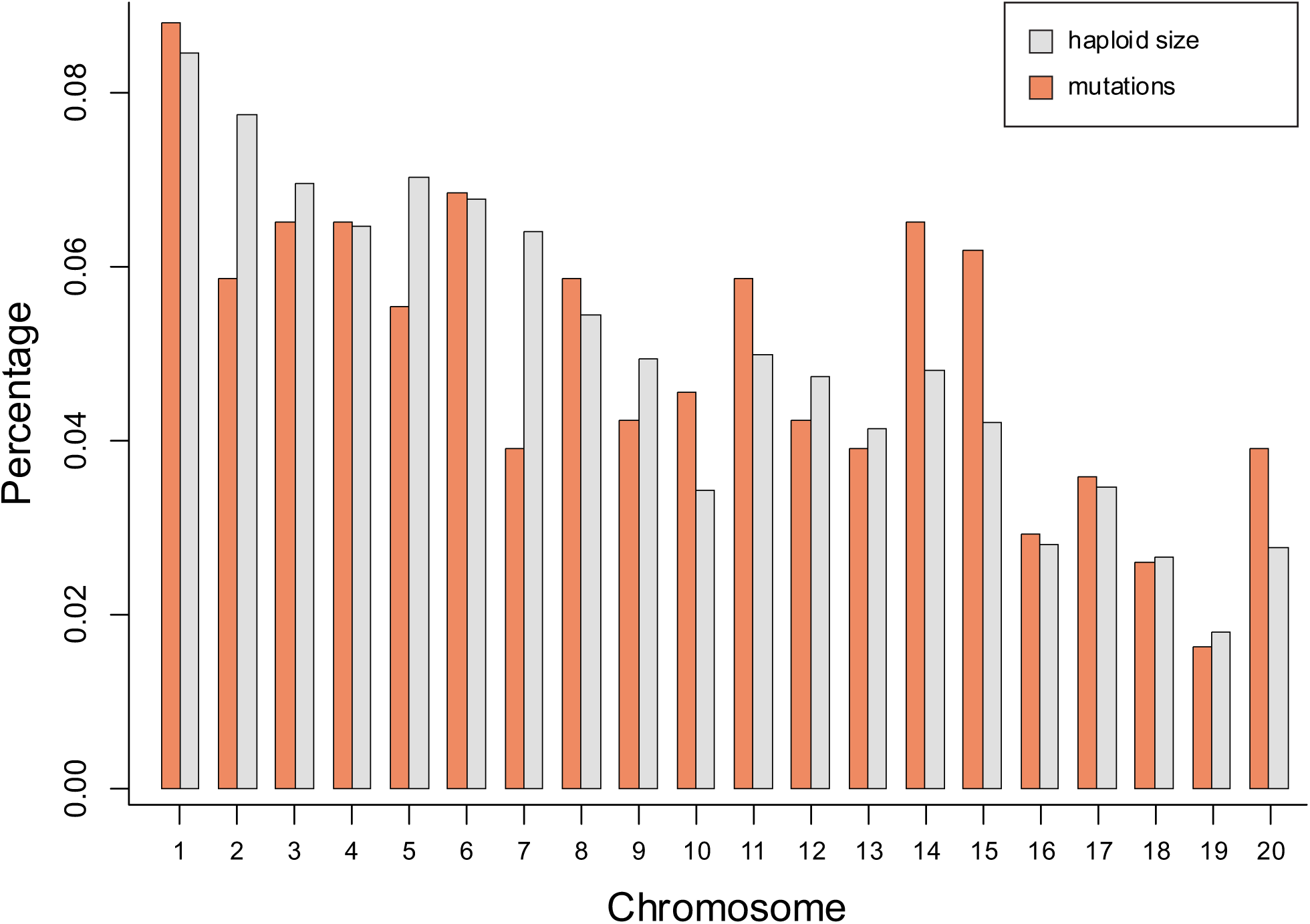
Percentage of *de novo* mutations identified on each autosome. The percentage of *de novo* mutations identified on each autosome, in orange, compared to the percentage of callable sites, in gray. Variation in frequency of mutations across chromosomes does not significantly differ from that expected from chance alone (χ^2^ test, *p* = 0.79).

**Figure S5.**
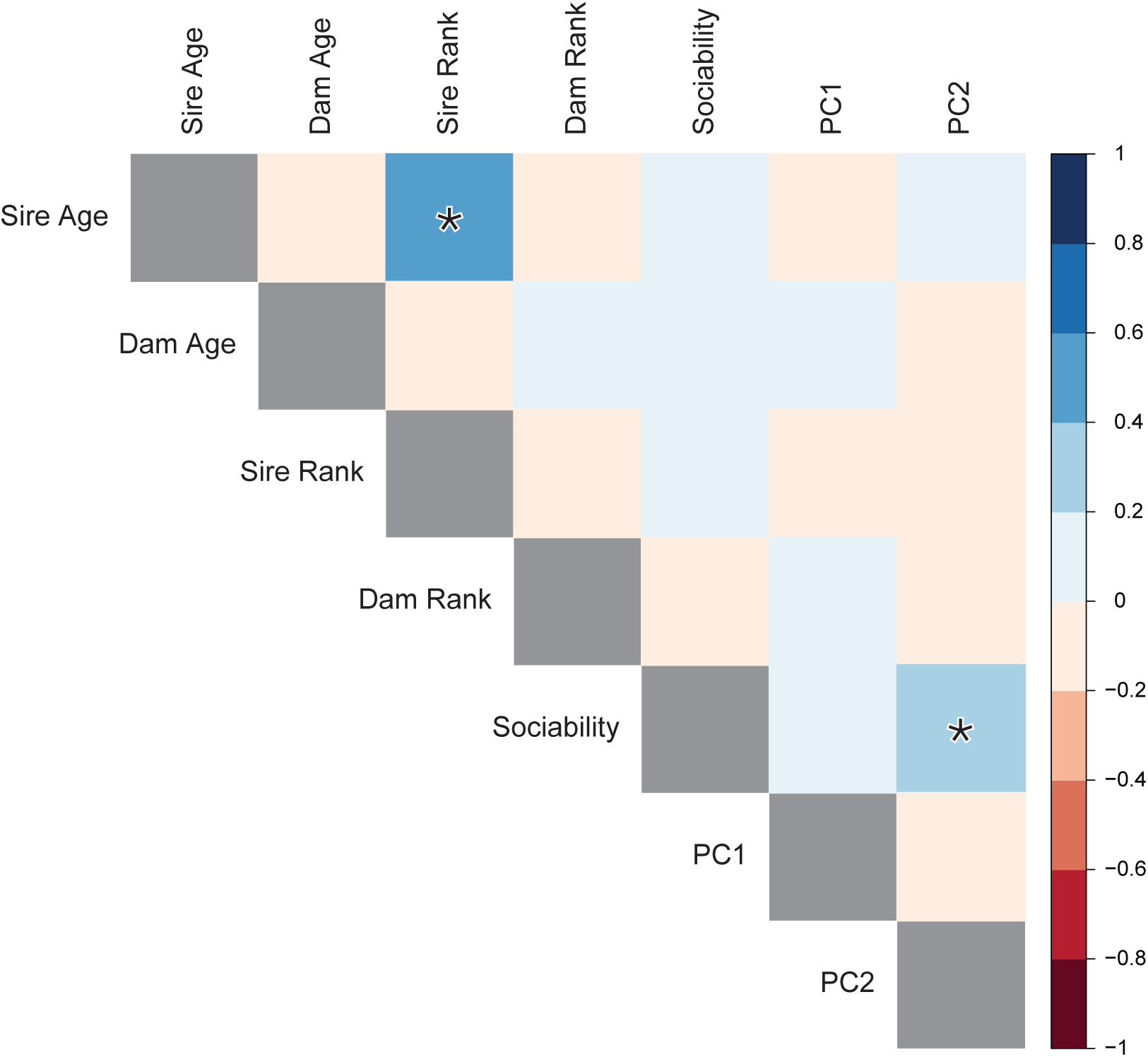
Correlations between parental age and behavioral traits in male rhesus monkeys. Boxes are shaded by the intensity of correlation in pairwise comparisons between each row and column. Diagonal values have been omitted. Legend on the right shows range of Pearson’s correlation coefficient for each color. Significant correlations (*p* < 0.05) are highlighted with an asterisk.

**Figure S6.**
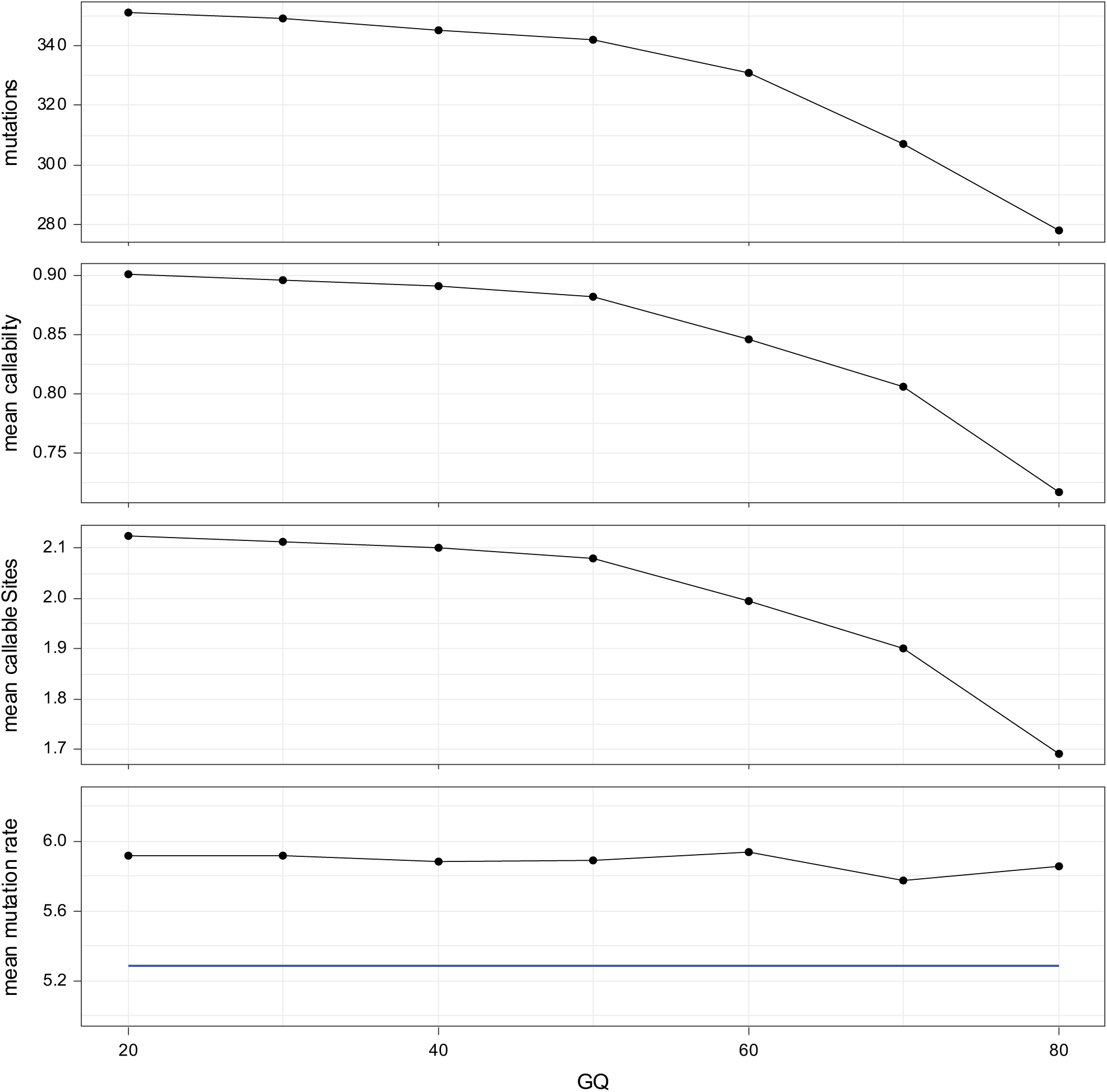
Filtering GATK candidate mutations by genotype quality. Plots show the effect of increasingly stringent GQ filters on the number of candidate mutations, mean callability, mean number of callable sites (×10^9^ bp), and the mean mutation rate (×10^-8^ per bp, per generation) across all trios. The mean mutation rate estimated from the *freebayes* calls is shown in blue for comparison.

**Figure S7.**
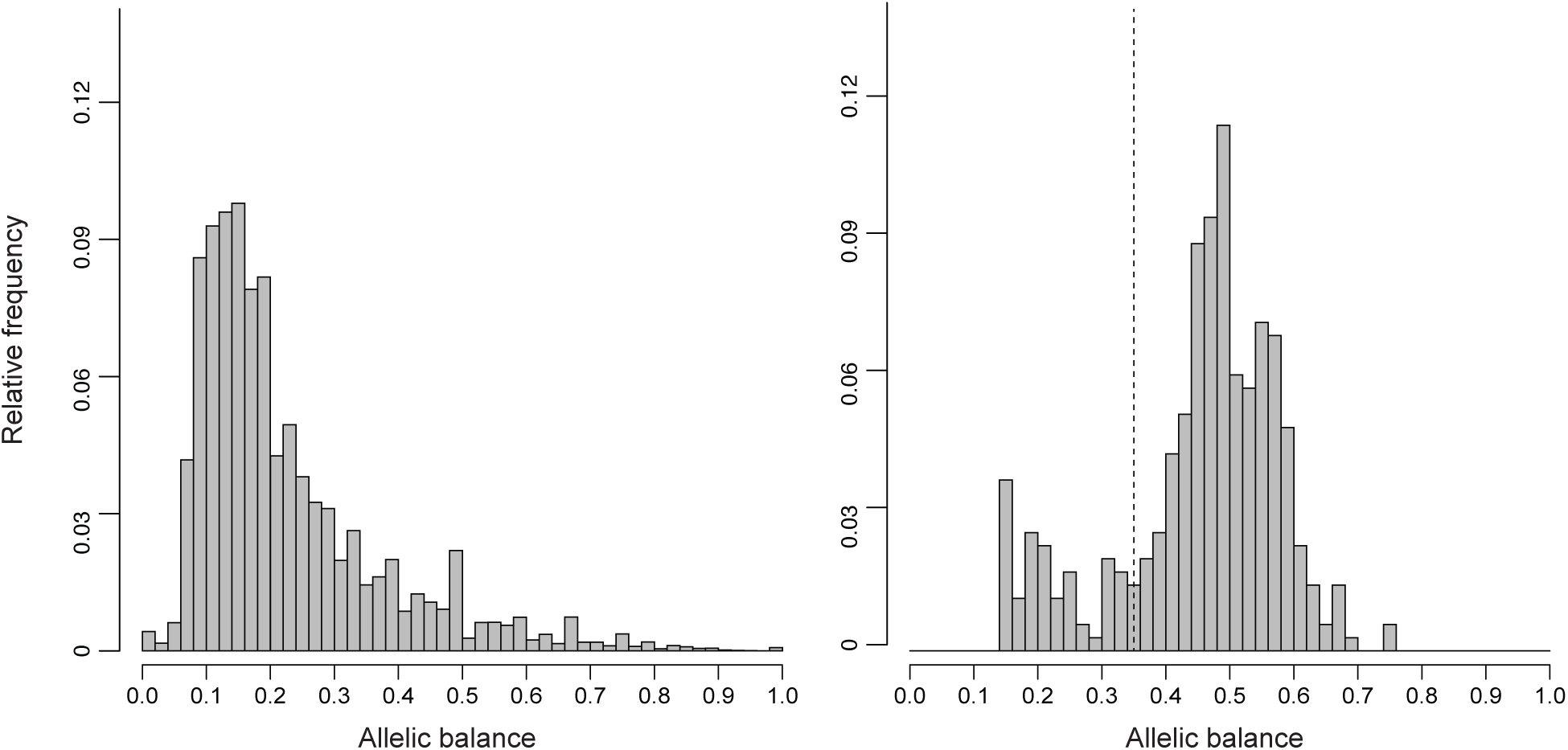
Allelic balance among candidate mutations. Distribution of allelic balance in heterozygote offspring that are Mendelian violations (left) and distribution at candidate *de novo* mutations after applying filters (right). Allelic balance at sites calculated as depth of alternate allele divided by total depth; dashed line indicates cutoff of 35% alternate depth.

**Table S1.** Loadings for first four principal components of behavioral observations

| Parameter<sup>a</sup> | PC1 | PC2 | PC3 | PC4 |
| --- | --- | --- | --- | --- |
| API(M) | -0.02 | 0.02 | -0.01 | -0.02 |
| API(F) | 0.02 | 0.01 | 0.03 | -0.02 |
| CON(M) | -0.30 | 0.49 | 0.01 | 0.65 |
| CON(F) | 0.44 | 0.31 | -0.81 | 0.10 |
| PRX(M) | -0.46 | 0.66 | 0.01 | -0.51 |
| PRX(F) | 0.70 | 0.46 | 0.54 | -0.04 |
| GRI(M) | -0.08 | 0.12 | -0.02 | 0.22 |
| GRI(F) | 0.07 | 0.04 | -0.20 | -0.51 |

<sup>a</sup> Abbreviations,
API approaches initiated to males(M) and females(F)
CON contact with males and females
PRX proximity to males and females
GRI grooming initiated to males and females

**Table S2.** Partial correlation of offspring sociability and sire age while controlling for sire rank

| Measure of social function | Pearson's *r* | *p*-value |
| --- | --- | --- |
| Sociability | 0.06 | 0.49 |
| Behavioral PC1 | -0.08 | 0.31 |
| Behavioral PC2 | 0.04 | 0.64 |
| Behavioral PC3 | -0.07 | 0.37 |
| Behavioral PC4 | 0.09 | 0.27 |

